# Nsp3 Ubl1-orchestrated dephosphorylation of N protein promotes coronaviral subgenomic RNA synthesis

**DOI:** 10.64898/2026.09.25.754410

**Authors:** Junji Zhu, Cindy Chiang, Michaela U. Gack

## Abstract

The coronavirus nucleocapsid (N) protein is indispensable for the viral lifecycle as part of the viral Replication-Transcription Complex together with Nsp3. Recent research demonstrated that phosphorylation of N by several host kinases intricately regulates its functions during infection. However, the mechanisms that control N dephosphorylation, and its physiological consequence, remain poorly understood. Here, we show that SARS-CoV-2 Nsp3 is a key orchestrator of N dephosphorylation by recruiting the phosphatase PP1α/γ via a conserved PP1-binding motif in the Ubl1 domain. Disruption of this motif abolishes N dephosphorylation and selectively impairs subgenomic RNA synthesis. Comparative interactome proteomics analysis of phospho-mimic vs. phospho-deficient N, together with functional validation, revealed that the host splicing factor SRSF1 cooperates with unphosphorylated N to promote subgenomic RNA transcription. These findings uncover an unrecognized mechanism in which Nsp3-guided N dephosphorylation by PP1α/γ enables coronavirus subgenomic RNA replication and highlight a potentially targetable axis for antiviral intervention.

## Introduction

Severe acute respiratory syndrome coronavirus (SARS-CoV-2) is a positive-sense single-stranded RNA virus that causes mild to severe respiratory disease and has sparked the COVID-19 global pandemic^1,2^. Despite the development of vaccines and antivirals, SARS-CoV-2 continues to circulate and cause acute disease of the respiratory tract or long-term pathology affecting multiple tissues (i.e., ‘long COVID’). SARS-CoV-2 encodes for ∼29 viral proteins –4 structural, 16 nonstructural (Nsps), and 9 accessory– that cooperatively regulate the specific steps of the viral life cycle from attachment and entry to viral release.

The nucleocapsid (N) protein is the most abundantly expressed protein of SARS-CoV-2 and possesses RNA binding and dimerization abilities that enable it to *1)* regulate viral RNA replication and transcription and *2)* package viral RNA within the ribonucleoprotein (RNP) complex^3–8^. How N protein is able to fulfill its distinct functions is still largely undetermined. However, a recent series of studies indicated that the phosphorylation state of N is key to regulating its specific activities inside infected cells. In the early stage of infection, phosphorylation of N at its serine-arginine-rich (SR) region by the kinases GSK-3, SRPK1/2, and CK1^9–12^ results in a conformational switch that reduces RNA-binding affinity, thereby facilitating the unpacking or release of viral RNA from N^13^. Furthermore, N undergoes liquid-liquid phase separation (LLPS) with viral RNA, forming condensates that facilitate virus replication^9–12,14–16^. Phosphorylation disrupts or fluidizes these N-RNA condensates, generating a more dynamic state compatible with transcriptional activity. By contrast, during the late stage of infection, N is predominantly unphosphorylated or hypophosphorylated within RNP complexes and virions, enabling tighter RNA binding that promotes RNP condensation and genome packaging^17^. Thus, N phosphorylation serves as a molecular switch that transitions the protein from an active regulator of viral RNA replication and transcription to a facilitator of virion formation. However, the exact functional consequences of N’s ‘phospho-switch’ and the phosphatase(s) that counteract N phosphorylation by GSK-3, SRPK1/2, and CK1 have not been determined.

Coronavirus replication takes place within double-membrane vesicles (DMVs), where the replication-transcription complex (RTC) is anchored to the inner membrane to provide a protected microenvironment for RNA synthesis^18^. Nsp3, the largest nonstructural protein of SARS-CoV-2, is the central scaffold of DMVs and serves as a multifunctional protein, aiding in polyprotein cleavage and processing, DMV formation, immune antagonism, and catalysis of de-ubiquitination and de-ISGylation events^18–23^. Among the multiple domains of Nsp3, the N-terminal ubiquitin-like 1 (Ubl1) domain of Nsp3 interacts with N as well as the 5′ UTR of the viral RNA and is pivotal for initiation of viral RNA synthesis and the enhancement of genomic RNA infectivity^24–29^. A recent study has shown that the Ubl1-N interaction disrupts LLPS of N and RNA^30^, suggesting that the Nsp3 Ubl1 domain may dynamically regulate N protein condensates to fine-tune RNA replication vs. packaging during different stages of the viral life cycle.

Coronavirus RNA synthesis proceeds through both continuous replication of full-length genomic RNA (gRNA) as well as discontinuous transcription of subgenomic mRNAs (sgmRNAs). The latter involves base-pairing between the nascent complementary body transcription regulatory sequence (cTRS-B) and the 5′ leader TRS (TRS-L), which mediates long-range template switching during minus-strand synthesis^31,32^. This leader-body fusion is directed by TRS complementarity and local RNA structures that transiently pause the polymerase, allowing template switching once a thermodynamic threshold is reached^33^. N protein plays a pivotal role in this process by recognizing TRS elements and modulating RNA conformation. Earlier studies showed that the N protein binds TRS-L with high affinity and recognizes TRS motifs across all sgmRNAs^34–36^. The N-terminal domain (NTD) of N protein then linearizes stem-loop 3 (SL3), which harbors the core TRS-L sequence, and melts TRS-cTRS duplexes, thereby exposing TRS-L for base-pairing with cTRS-B to ultimately promote template switching^37^. Phosphorylated N also recruits the RNA helicase DDX1 to enhance TRS-B readthrough, facilitating synthesis of full-length gRNA and longer sgmRNAs^38^. Given that in vitro template-switching assays typically employ bacterially expressed, unphosphorylated N protein and since phosphorylation reduces RNA-binding affinity, it is likely that the unphosphorylated form of N protein predominantly supports efficient template switching and discontinuous transcription. However, the molecular mechanism(s) by which N transitions between phosphorylated and unphosphorylated states during the viral replication cycle, and how the dynamic balance of phospho-vs. non-phospho N governs discontinuous transcription in infected cells, remain unresolved.

In this study, we identified that SARS-CoV-2 Nsp3 recruits the host phosphatase PP1α/γ via a conserved PP1-binding motif in the Ubl1 domain, thereby facilitating N dephosphorylation. Proteomics analysis combined with molecular characterization studies further demonstrated that N dephosphorylation by PP1 is essential for SARS-CoV-2 subgenomic RNA synthesis and mediated by the splicing factor SRSF1.

## Results

### SARS-CoV-2 Nsp3 induces N dephosphorylation in a PP1α/γ-dependent manner

SARS-CoV-2 N is a modular RNA-binding protein in which the NTD and C-terminal domain (CTD) are embedded within a flexible architecture of intrinsically disordered regions (IDR), including the N-IDR, the serine/arginine-rich linker region (LKR), and the C-IDR (Fig. 1a). The CTD contributes to both RNA engagement and N self-association, whereas the SR-rich LKR is a major regulatory hub for phosphorylation-dependent control of N activity. To examine how N phosphorylation influences its interaction with the viral replication machinery, we generated two previously characterized phosphorylation-state mutants: N-2SA, in which the GSK-3 priming sites S188 and S206 were replaced with alanines, and N-10D, in which ten serine/threonine residues within the SR-rich region were substituted with aspartates to mimic constitutive phosphorylation^39^ (Fig. 1a). Given that N phosphorylation has been linked to distinct stages of viral RNA synthesis and RNP assembly^38–40^, and since Nsp3-Ubl1 is known to recruit N to DMV-associated replication sites^4,25^, we asked whether Nsp3 binding is influenced by the phosphorylation state of N.

**Fig. 1.**
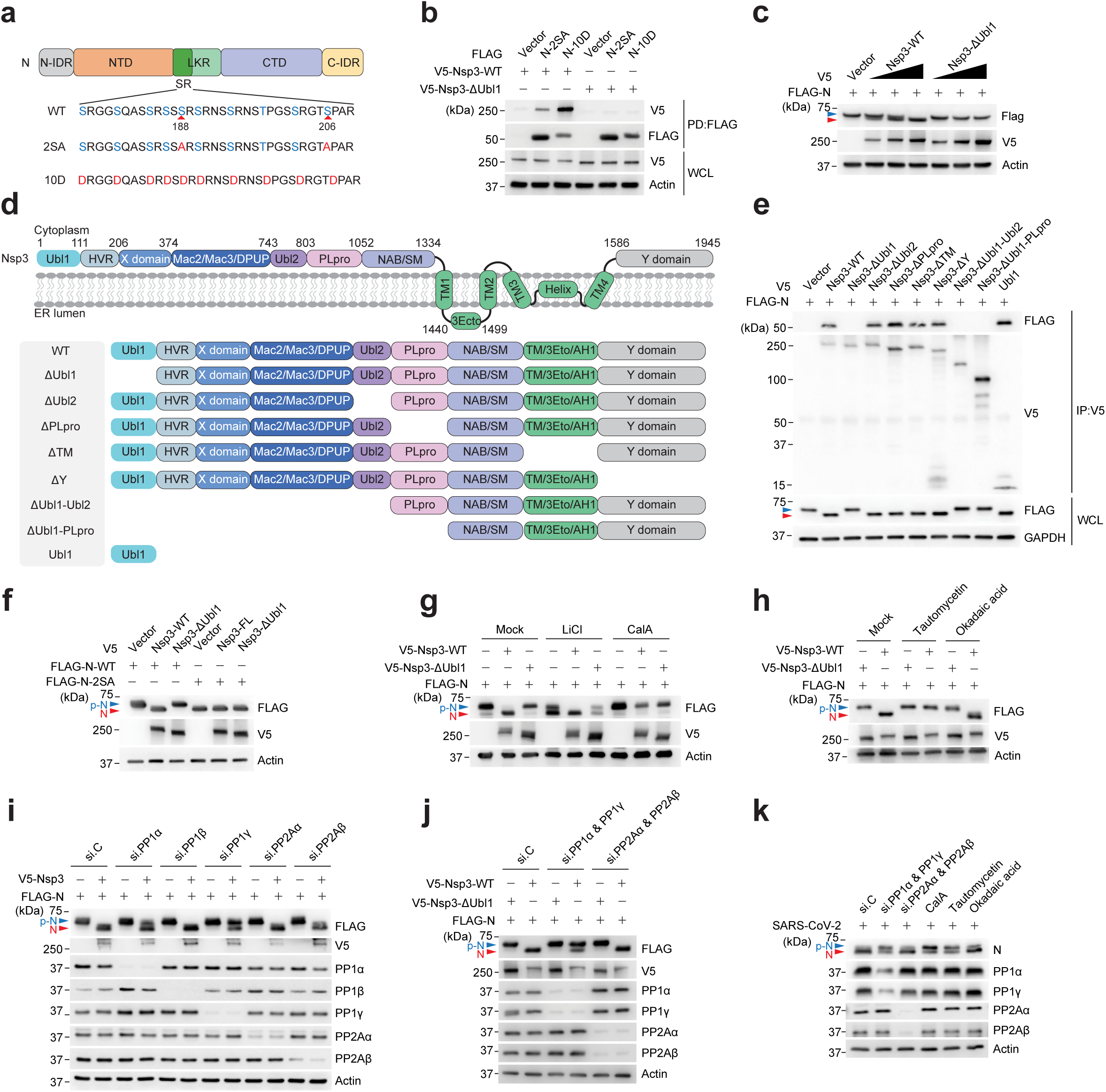
SARS-CoV-2 Nsp3 promotes N dephosphorylation in a PP1α/γ-dependent manner. **a**, Schematic of the SARS-CoV-2 N protein’s domain architecture (upper) and its serine/arginine-rich (SR) region (lower), with key serine residues and the priming phosphorylation sites S188 and S206 highlighted. The specific mutations in the phospho-deficient (2SA) and phosphomimetic (10D) mutants are indicated in red. **b**, Nsp3 interaction of N-2SA or N-10D in HEK293T cells that were transfected for 24 h with either empty vector (−) or FLAG-tagged N-S2A or N-10D along with V5-tagged Nsp3 WT or ΔUbl1 mutant, determined by PD:FLAG and immunoblot (IB) with anti-V5. **c**, IB analysis of the canonical and faster-migrating forms of FLAG-tagged N protein in the whole cell lysates (WCLs) of HEK293T cells that were transfected for 24 h with either empty vector (−) or V5-tagged Nsp3 WT or ΔUbl1 mutant. **d**, Upper: Schematic diagram of SARS-CoV-2 Nsp3’s domain architecture and topology. Numbers indicate amino acids. Lower: Nsp3 WT and generated truncation mutants. **e**, Binding of FLAG-tagged N protein to Nsp3 in HEK293T cells that were transiently transfected with either empty vector (−) or V5-tagged full-length Nsp3 (Nsp3 WT) or the indicated mutants, determined by IP:V5 and IB with anti-FLAG. The canonical and faster-migrating forms of FLAG-tagged N protein were determined in the WCLs by IB with anti-FLAG. **f**, IB analysis showing the electrophoretic mobility of FLAG-tagged N WT or 2SA mutant in WCLs of HEK293T that were co-transfected for 24 h with either vector (−) or V5-tagged Nsp3 WT or ΔUbl1 mutant. **g**, IB analysis of N protein phosphorylation status in the WCLs of HEK293T cells that were either mock-treated or pretreated with 50 mM lithium chloride (LiCl) or 100 nM Calyculin A (CalA) and transfected for 24 h with FLAG-tagged N along with either empty vector (−) or V5-tagged Nsp3 WT or ΔUbl1. **h**, IB analysis of FLAG-tagged N protein phosphorylation status in the WCLs of HEK293T cells that were either mock treated or pretreated with 5 µM Tautomycetin or 100 nM Okadaic acid and transfected for 24 h with either empty vector (−) or V5-tagged Nsp3 WT or ΔUbl1. **i, j**, IB analysis of FLAG-tagged N protein phosphorylation status in the WCLs of HEK293T cells that were transfected for 48 h with the indicated siRNAs and then transfected for 24 h with either empty vector (−) or V5-tagged Nsp3 WT or ΔUbl1. **‘**si.C’ indicates nontargeting control siRNA. **k**, IB analysis of N protein phosphorylation status in the WCLs of A549-ACE2 cells that were transfected for 48 h with the indicated siRNAs and then infected with SARS-CoV-2 (strain K49; MOI 1) for 12 h. Blue and red arrows indicate the phosphorylated and unphosphorylated N protein forms respectively. Data are representative of at least two (**b**-**c, e**-**k**) independent experiments.

Co-immunoprecipitation (Co-IP) analysis showed that wild-type (WT) Nsp3 associated more efficiently with N-10D (phosphomimetic) than with N-2SA (phospho-‘null’). This preferential binding was lost when the N-terminal Ubl1 domain of Nsp3 was deleted (ΔUbl1), indicating that Ubl1 mediates N binding while conferring preferential recognition of phosphorylated N (Fig. 1b). We also noted that Nsp3 co-expression caused N to migrate faster by SDS-PAGE, and this shift was induced by WT Nsp3 but not by Nsp3-ΔUbl1, linking the altered electrophoretic mobility of N to its Ubl1-dependent interaction with Nsp3 (Fig. 1c).

To further define the Nsp3 region responsible for these effects, we generated a panel of Nsp3 truncation constructs covering its major cytosolic and membrane-associated segments (Fig. 1d). Only Nsp3 fragments that retained Ubl1 bound to N and induced its electrophoretic mobility shift, whereas fragments lacking Ubl1 did not (Fig. 1e). Notably, expression of Ubl1 alone was sufficient to reproduce this phenotype (Fig. 1e). Thus, the Ubl1 domain is not merely required for Nsp3-N binding, but constitutes the minimal Nsp3 element required to trigger the mobility change of N.

Importantly, the faster-migrating band for N induced by Nsp3 Ubl1 was observed for WT N but not for phospho-deficient N-2SA, which already exhibits increased electrophoretic mobility due to the loss of phosphorylation^10^ (Fig. 1f). This suggests that the shift induced by Nsp3 Ubl1 likely reflects reduced N phosphorylation (Fig. 1f). Consistent with this proposal, the Nsp3-induced mobility shift of N phenocopied the effect of N phosphorylation inhibition by lithium chloride (LiCl), an inhibitor of GSK-3 which is a key kinase mediating coronavirus N phosphorylation^10^. By contrast, treatment of cells with calyculin A (Cal A), a highly specific inhibitor of the serine/threonine protein phosphatases 1 (PP1) and 2A (PP2A), abolished this effect (Fig. 1g). These observations suggest that Nsp3 promotes N dephosphorylation rather than preventing its phosphorylation.

We next sought to identify the phosphatase(s) responsible for N dephosphorylation and ‘mobility shift’. Pharmacological inhibition with tautomycetin, which preferentially targets PP1, prevented the Nsp3-induced N mobility shift, whereas okadaic acid, used under conditions favoring PP2A inhibition^41^, had little effect (Fig. 1h). In accord, siRNA-mediated depletion of PP1α and/or PP1γ impaired Nsp3-driven N dephosphorylation, whereas depletion of PP1β or of the PP2A isoforms PP2Aα and PP2Aβ had no such effect (Fig. 1i,j). Similarly, during authentic SARS-CoV-2 infection, PP1α/γ depletion, or treatment with CalA or tautomycetin, increased the phosphorylated N protein pool, whereas PP2Aα/β depletion or okadaic acid treatment had a minimal impact (Fig. 1k).

These data identify the Ubl1 domain of Nsp3 as a key determinant that recognizes phosphorylated N and promotes its dephosphorylation by PP1α/γ. This mechanism provides a direct link between N recruitment to replication organelles and the dynamic regulation of the N phosphorylation state during SARS-CoV-2 infection.

### PP1α/γ recruitment to the Nsp3 Ubl1 domain via a conserved PP1-binding motif leads to N dephosphorylation

We next investigated whether Nsp3 physically recruits PP1α and PP1γ. Co-IP analysis showed that WT Nsp3 efficiently interacted with PP1α and PP1γ, but minimally with PP1β, whereas deletion of the Ubl1 domain abolished these interactions (Fig. 2a). Consistently, WT Nsp3, but not Nsp3-ΔUbl1, interacted with endogenous PP1α and PP1γ, while no detectable association was observed with PP1β, PP2Aα, or PP2Aβ (Fig. 2b). These data indicated that the Ubl1 domain is required for Nsp3 association with the PP1α/γ isoforms implicated in N dephosphorylation.

**Fig. 2.**
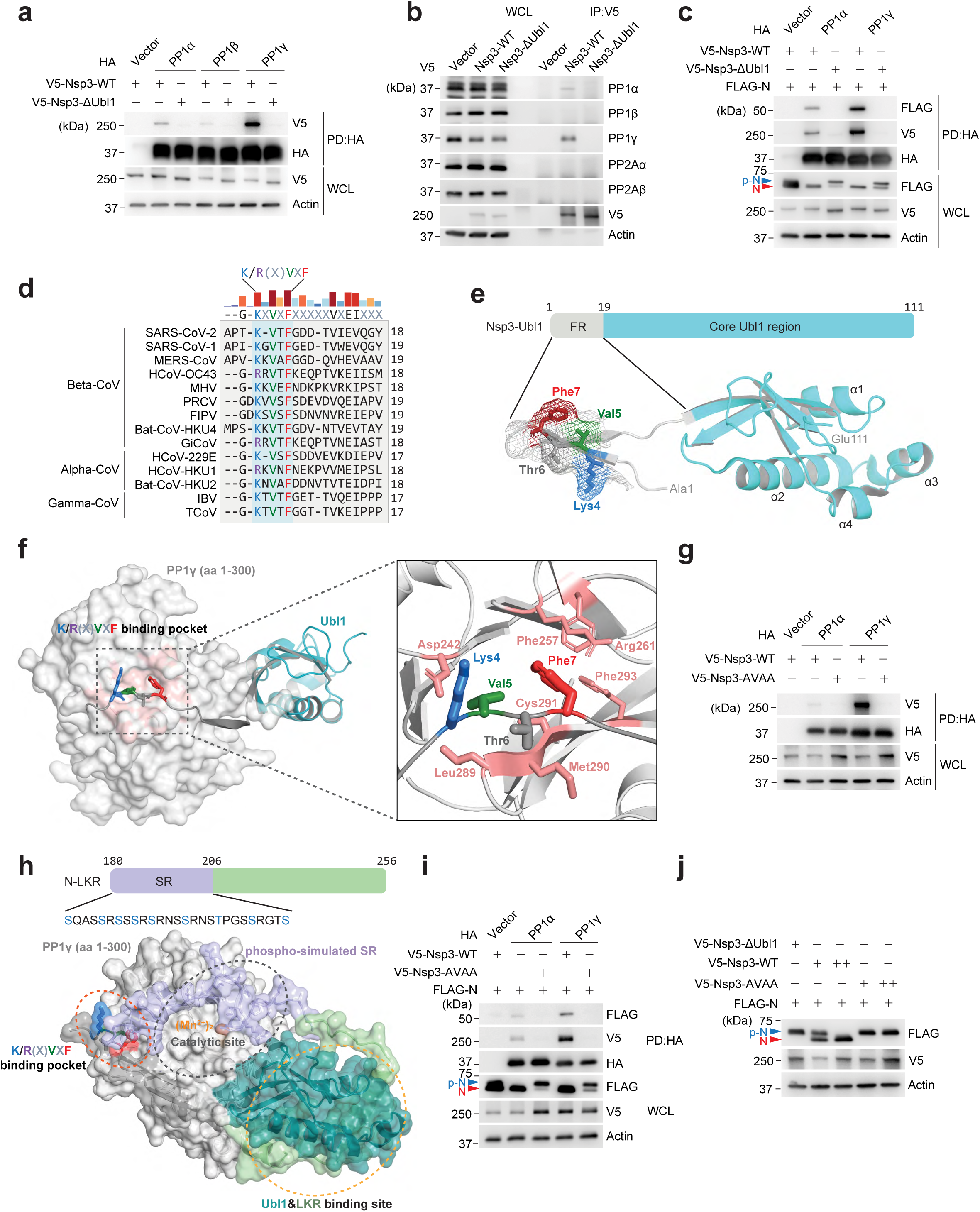
PP1α/γ recruitment to the Nsp3 Ubl1 domain via a conserved PP1-binding motif leads to N dephosphorylation. **a**, Nsp3 binding of HA-tagged PP1α, PP1β or PP1γ in HEK293T cells that were co-transfected for 24 h with V5-tagged Nsp3 WT or ΔUbl1, determined by HA pulldown (PD:HA) and IB with anti-HA. **b**, Binding of Nsp3 to endogenous PP1 or PP2 proteins in HEK293T cells that were transfected for 24 h with V5-tagged Nsp3 WT or ΔUbl1, determined by IP:V5 and IB with indicated antibodies. **c**, N binding of HA-tagged PP1α or PP1γ in HEK293T cells that were transfected for 24 h with FLAG-tagged N together with either V5-tagged Nsp3 WT or ΔUbl1, determined by PD:HA and IB with anti-FLAG, anti-V5, and anti-HA. **d**, Sequence alignment of the Nsp3 Ubl1 domain from the indicated alpha-, beta-, and gamma-coronaviruses reveals a conserved PP1-binding motif K/R(X)VXF. Numbers indicate amino acids. **e**, Upper: Schematic diagram of the Nsp3 Ubl1 domain containing a flexible region (FR) and a core Ubl1 region. Lower: Representation of the Nsp3 Ubl1 structure (PDB: 7KAG) with the PP1-binding motif residues highlighted. **f**, AlphaFold3 prediction of Nsp3 Ubl1 and PP1γ binding (pTM: 0.85; ipTM: 0.83), with the KVTF residues in Nsp3 Ubl1 highlighted in red, green, blue, and grey, and the PP1γ residues forming the binding pocket highlighted in pink. **g**, PP1α/γ binding of Nsp3 in HEK293T cells that were transfected for 24 h with either empty vector (−) or HA-tagged PP1α or PP1γ together with V5-tagged Nsp3 WT or AVAA mutant, determined by PD:HA and IB with anti-V5. **h**, Upper: Schematic diagram showing the N protein linker region (N-LKR), with the indicated residues in the SR region simulated to phosphorylated Ser or Thr. Lower: AlphaFold3 prediction of Nsp3 Ubl1, phospho-simulated N-LKR and PP1γ (aa 1-300) binding (pTM: 0.77; ipTM: 0.74). The PP1-binding pocket, catalytic site, and the Ubl1-LKR binding site are highlighted. **i**, PP1α/γ binding of FLAG-tagged N protein in HEK293T cells that were transfected for 24 h with either V5-tagged Nsp3 WT or AVAA mutant together with either empty vector (−) or HA-tagged PP1α or PP1γ, determined by PD:HA and IB with anti-FLAG, anti-V5, and anti-HA. **j**, IB analysis of FLAG-tagged N protein in HEK293T cells that were co-transfected for 24 h with either V5-tagged Nsp3 WT, ΔUbl1, or AVAA mutant. Blue and red arrows indicate the phosphorylated and unphosphorylated N protein forms respectively. Data are representative of at least two (**a**-**c, g, i** and **j**) independent experiments.

Consistent with this concept, Co-IP analysis showed that PP1α and PP1γ associated with N after co-expression of WT Nsp3 but not Nsp3-ΔUbl1. Furthermore, in the absence of WT Nsp3, PP1α/γ was not sufficient to effectively dephosphorylate N, suggesting that Nsp3 and specifically its Ubl1 region is required to bridge PP1α/γ and N for promoting N dephosphorylation (Fig. 2c). Prompted by this observation, we inspected the SARS-CoV-2 Ubl1 sequence and found a short K/R(X)VXF-like sequence, ^4^KVTF^7^, resembling the canonical PP1-docking motif used by many PP1-regulatory proteins. This motif was highly conserved across representative alpha-, beta-, and gamma-coronaviruses (Fig. 2d). The KVTF motif locates to the flexible N-terminal region of Ubl1, which is upstream of the structured Ubl1 core (Fig. 2e). Consistent with its predicted function as a PP1-docking element, AlphaFold3 modelling positioned the KVTF motif within the conserved docking groove of PP1γ, with the Val and Phe residues occupying the hydrophobic pocket typically engaged by PP1-interacting proteins (Fig. 2f).

To test whether this motif mediates direct PP1 recruitment, we replaced KVTF with AVAA. This mutation nearly abolished Nsp3 binding to both PP1α and PP1γ (Fig. 2g), confirming that this conserved PP1-binding motif is required for efficient PP1 recruitment. Given that the N-LKR domain contains both the phosphoregulatory SR and two linear motifs that mediate binding to the Nsp3 Ubl1^24^, we next modelled using Alphafold3 a ternary complex containing Nsp3 Ubl1, PP1γ (aa 1-300), and an N-LKR peptide carrying phospho-simulated SR (Fig. 2h). In this model, the folded core of Ubl1 engages the N-LKR segment through the two binding motifs as described previously^24^, while the Ubl1 KVTF sequence docks into the canonical K/RVxF-binding pocket of PP1γ. This ternary arrangement tethers PP1γ to the Ubl1-N-LKR complex and places the phosphorylated SR of N-LKR within reach of the PP1γ catalytic site (Fig. 2h). By contrast, models containing unmodified N-LKR placed the SR farther from the catalytic center (Extended Data Fig. 1a,b). In accord with this model, the Nsp3 AVAA mutant failed to promote PP1-N complex formation and efficient N dephosphorylation despite retaining N-binding activity (Fig. 2i, j and Extended Data Fig. 1c).

Together, these data identify a conserved PP1-binding motif in the Ubl1 domain of coronaviral Nsp3 that is required for PP1α/γ recruitment and PP1α/γ-mediated N dephosphorylation. Given that N is predominantly unphosphorylated within RNP complexes and virions, these findings support a model in which Nsp3, besides mediating the export of gRNA, orchestrates N dephosphorylation to promote efficient RNP packaging at the DMV during the late phase of infection.

### Nsp3-directed N dephosphorylation is required for subgenomic RNA synthesis

During the early phase of infection, N is actively involved in viral RNA replication^4,38,42^. To examine the functional consequence of Nsp3-orchestrated N dephosphorylation in viral RNA replication, we used a single-cycle SARS-CoV-2 replicon in which the spike coding sequence is replaced by an mScarlet reporter, allowing reporter expression to serve as a readout of subgenomic RNA production^43^. As mutation of the conserved Phe (F→A) within the Ubl1 ^4^KVTF^7^ motif was sufficient to impair PP1 binding and prevent Nsp3-guided N dephosphorylation by PP1, similarly to the AVAA mutant (Extended Data Fig. 2a,b), we introduced the F^7^→A^7^ substitution into the replicon, generating Rep.KVTA (Fig. 3a). Co-IP showed that WT replicon (Rep.WT)-derived Nsp3 associated with endogenous PP1α and PP1γ, whereas Nsp3 encoding the KVTA mutation did not bind PP1α/γ (Fig. 3b). Consistent with defective PP1 recruitment, Rep.KVTA produced only phosphorylated N, in contrast to Rep.WT, in which both phosphorylated and unphosphorylated N were detected (Fig. 3c). These results confirm that N dephosphorylation requires PP1 recruitment to Nsp3-Ubl1 during SARS-CoV-2 replication.

**Fig. 3.**
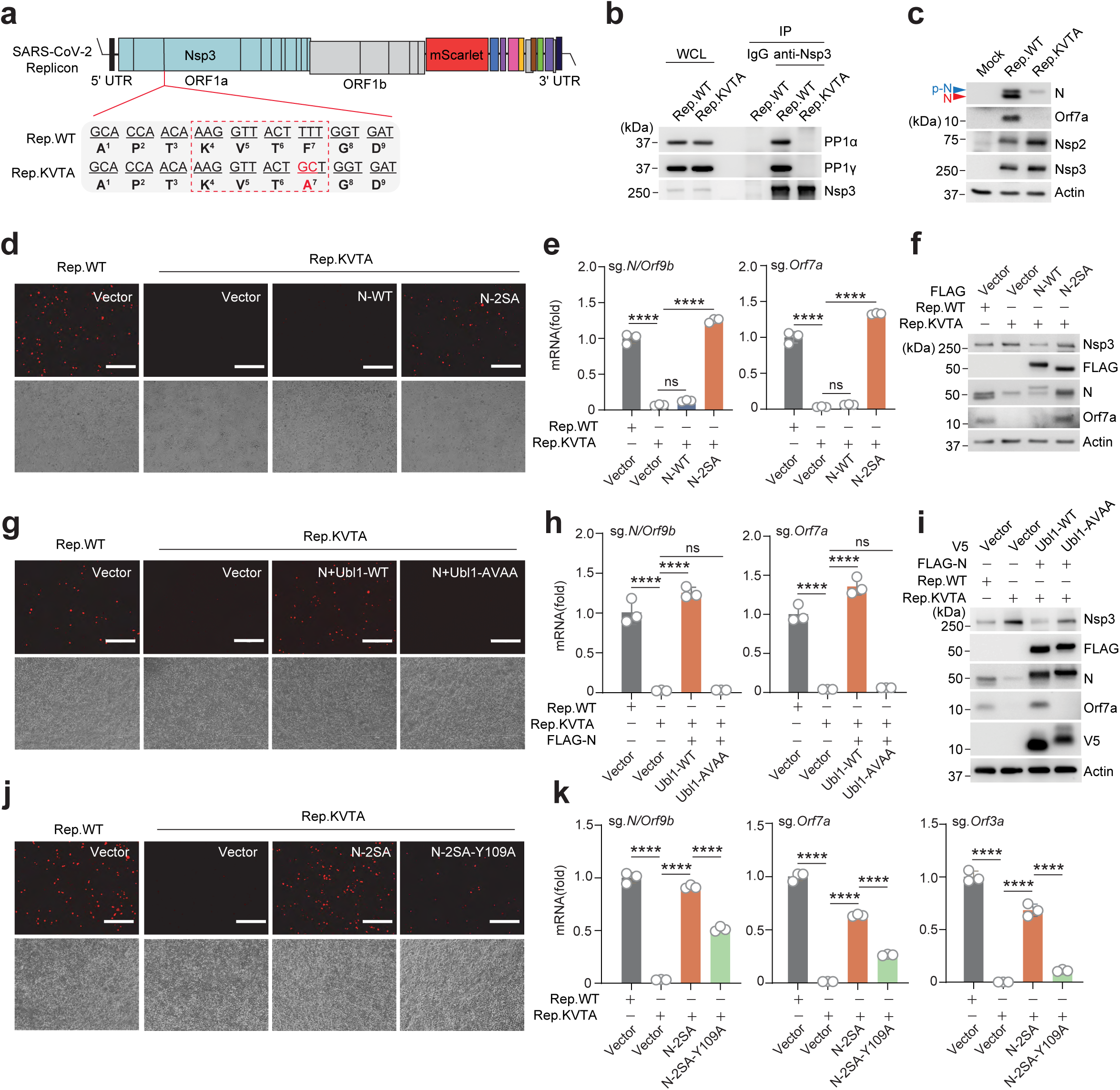
N dephosphorylation is required for viral subgenomic RNA synthesis. **a**, Schematic diagram of the SARS-CoV-2 replicon (Rep.WT) in which the spike gene was replaced with mScarlet, as well as of the mutant replicon (Rep.KVTA) in which Phe7 in Nsp3 was mutated to Ala7 (red). **b**, PP1α/γ and N binding to Nsp3 in HEK293T cells that were transfected for 48 h with either Rep.WT or Rep.KVTA, determined by IP:Nsp3 and IB with the indicated antibodies. **c**, IB analysis of the indicated SARS-CoV-2 proteins in HEK293T cells that were either mock treated or transfected for 48 h with either Rep.WT or Rep.KVTA. **d**-**f**, Replication of Rep.WT or Rep.KVTA in HEK293T cells that were transfected for 48 h with Rep.WT or Rep.KVTA and either empty vector, FLAG-tagged N WT or 2SA mutant, determined by immunofluorescence (IF) microscopy analysis of mScarlet signals (**d**), RT-qPCR analysis of the indicated viral sg.RNAs (**e**) and IB analysis of the indicated SARS-CoV-2 proteins (**f**). **g**-**i**, Replication of Rep.WT or Rep.KVTA in HEK293T cells that were transfected for 48 h with Rep.WT or Rep.KVTA and either empty vector or FLAG-N together with V5-tagged Nsp3-Ubl1 WT or AVAA, determined as in (d-f). **j**,**k**, Replication of Rep.WT or Rep.KVTA in HEK293T cells that were transfected for 48 h with Rep.WT or Rep.KVTA and either empty vector or FLAG-tagged N-2SA or N-2SA-Y109A, determined by IF microscopy analysis of mScarlet signals (**j**) or RT-qPCR analysis of the indicated viral sg.RNAs (**k**). Data are representative of at least two (**b**-**k**) independent experiments. ****P < 0.0001 (one-way ANOVA with Sidak’s multiple comparison test in **e**, **h** and **k**). ns, not significant.

Notably, mScarlet mRNA and viral subgenomic RNAs were markedly reduced in a position-dependent manner, with stronger defects for transcripts located farther from the 3′ UTR: ∼50-fold for sg.*ORF3a*, ∼20-fold for sg.*ORF7a* and ∼10-fold for sg.*N/ORF9b.* In line with this, N and ORF7a protein expressions were also decreased (Fig. 3c and Extended Data Fig. 2c). However, Rep.KVTA showed increased genomic RNA levels (e.g., g.*Nsp3*) and higher abundance of several nonstructural proteins including Nsp2 and Nsp3 (Fig. 3c and Extended Data Fig. 2c). These results suggest that disruption of the PP1-docking motif in Nsp3 selectively impairs subgenomic RNA synthesis (rather than broadly suppressing replicon amplification), which is likely caused by dysregulated discontinuous transcription.

We next asked whether this defect was attributed to impaired N dephosphorylation. Fluorescence microscopy analysis showed that Rep.KVTA exhibited diminished mScarlet signal compared with Rep.WT (Extended Data Fig. 2d,e). This defect was partially restored by ectopic expression of WT Nsp3, either alone or together with N (Extended Data Fig. 2d,e). Crucially, expression of the phospho-deficient N-2SA mutant fully restored mScarlet signal to WT levels, while the expression of WT N had only a small effect (Fig. 3d and Extended Data Fig. 2d,e). RT-qPCR analysis further showed that N-2SA, but not WT N, restored the abundance of Rep.KVTA-derived subgenomic RNAs, including sg.*N/Orf9b* and sg.*Orf7a* (Fig. 3e), as well as ORF7a protein expression (Fig. 3f). By contrast, the phosphomimetic N-10D mutant failed to restore Rep.KVTA mScarlet signal or ORF7a expression (Extended Data Fig. 2f,g). Consistently, WT Ubl1, but not Ubl1-AVAA, restored Rep.KVTA mScarlet signal (Fig.3g and Extended Data Fig. 2d,e), sg.*N/Orf9b* and sg.*Orf7a* RNA levels (Fig. 3h), as well as ORF7a protein expression (Fig. 3i).

Given that N modulates viral RNA replication through its RNA chaperone activity, and since Y109 within the NTD of N is critical for viral RNA binding^44–47^, we next introduced the Y109A substitution into the N-2SA mutant. Compared with N-2SA, N-2SA-Y109A showed a markedly impaired rescue activity in Rep.KVTA (Fig. 3j and Extended Data Fig. 2h). RT-qPCR analysis confirmed that, as compared to N-2SA, N-2SA-Y109A was less effective in restoring subgenomic RNAs, including sg.*N/Orf9b*, sg.*Orf7a* and sg.*Orf3a*, showing an indirect correlation between rescue efficiency and transcript distance from the 3′ UTR (Fig. 3k). This suggests that N-2SA-mediated rescue, at least in part, requires the RNA-binding function of N.

Together, these data support a model in which Nsp3-mediated PP1 recruitment generates a dephosphorylated, RNA-binding-competent N protein pool for efficient subgenomic RNA synthesis.

### The phosphorylation state of N determines its host interactome landscape

As the phospho-deficient N mutant but not the phosphorylated N, restored subgenomic RNA synthesis of Rep.KVTA, and because it has been shown that specific host RNA-binding proteins (RBPs) engage SARS-CoV-2 RNA and N to regulate RNA replication^48–53^, we next asked whether N phosphorylation shapes its host-interacting protein network. To this end, FLAG-tagged N-2SA (phospho-ablated) and N-10D (phosphomimetic) expressed in mammalian cells were subjected to FLAG affinity purification followed by LC-MS/MS analysis of N-associated proteins (Fig. 4a). Comparative interactome analysis identified 146 proteins preferentially enriched with N-2SA, and 176 proteins preferentially enriched with N-10D. Gene ontology analysis revealed partially overlapping but functionally distinct enrichments between the two datasets (Fig. 4b,c). Proteins that were preferentially associated with N-2SA were enriched primarily for mRNA processing (Fig. 4b). By contrast, the N-10D-enriched interactome showed an enrichment for translation and ribosome biogenesis (Fig. 4c). In support of this, KEGG pathway analysis showed that N-2SA-associated proteins were mainly enriched in spliceosome and mRNA surveillance pathways, whereas N-10D-associated proteins were predominantly involved in ribosome-related pathways and protein groups linked to coronavirus disease (Fig. 4d).

**Fig. 4.**
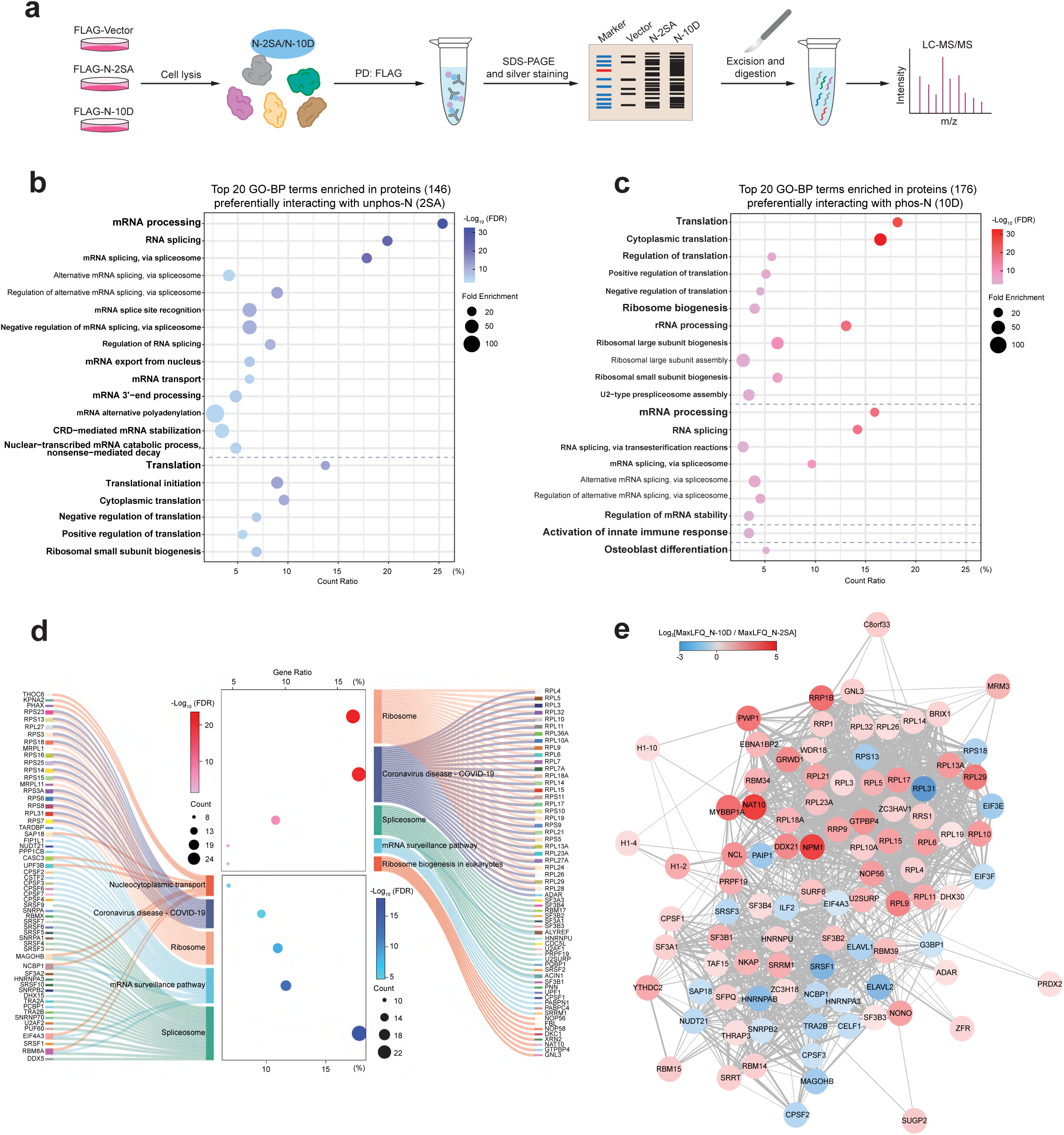
Comparative proteomics analysis identifies distinct interactomes of phosphorylated and unphosphorylated N. **a**, Schematic overview of the workflow for the affinity purification-mass spectrometry (AP-MS) analysis to identify distinct interactomes of phospho-null (N-S2A) vs phosphomimetic (N-10D) N protein. HEK293T cells were transfected with either FLAG-tagged N-2SA or N-10D. 48 h later, N protein was purified from total cell lysates by PD:FLAG, followed by SDS-PAGE and LC-MS/MS analysis of co-purified proteins (see also Methods). **b**,**c**, Dot plots of GO-BP enrichment analysis showing the top twenty biological processes associated with the N-2SA interactome (146 proteins) (**b**) or N-10D interactome (176 proteins) (**c**). Related GO-BP terms were grouped according to their broader parent biological-process categories, with horizontal dashed lines indicating the boundaries between these groups. Dot color reflects FDR-adjusted p value, and dot size represents fold enrichment. **d**, KEGG pathway analysis showing the enriched top five pathways for the N-2SA (146 proteins) and N-10D (176 proteins) interactome. Left and right: Sankey plot illustrating the mapping of interactome of N-2SA (left) and N-10D (right) to their associated KEGG pathways; Central: dot plot showing significantly enriched KEGG pathways of N-10D (up) and N-2SA (down), ranked by gene ratio and color-coded by –log_10_(FDR). Dot size represents the number of genes per pathway. **e**, Visualization of the comparative protein-protein interactome of N-2SA and N-10D [|log_2_(MaxLFQ_10D/MaxLFQ_2SA)| > 1]. Protein-protein interactions were retrieved from the STRING database and visualized using Cytoscape. Edges represent known or predicted protein-protein interactions from STRING, and color gradient reflects the log_2_ fold change of N-10D / N-2SA MaxLFQ intensities. Data shown are from one unbiased AP-MS screen.

To visualize how these phosphorylation-state-dependent interactions were organized, we generated Cytoscape networks of proteins differentially enriched with N-2SA vs. N-10D. This analysis revealed two separable host factor modules (Fig. 4e and Extended Data Fig. 3a,b). The N-2SA-enriched network contained multiple RNA-binding and RNA-processing factors, including SRSF-family proteins, HNRNP proteins, and other components implicated in RNA maturation or trafficking. By contrast, the N-10D-enriched network was dominated by ribosomal proteins and translation-associated factors, particularly members of the ribosomal protein large subunit (RPL) family. Thus, these data show that the phosphorylation state of N is a major determinant of its host-interactome landscape.

### SRSF1 is required for dephosphorylated N-mediated subgenomic RNA synthesis

We next sought to identify host factors that cooperate with dephosphorylated N to mediate viral subgenomic RNA synthesis. From the N-2SA interactome, we selected candidate proteins that were enriched by more than 2-fold in MaxLFQ intensity compared to N-10D for a targeted siRNA screen to assess their effects on viral RNA replication using the SARS-CoV-2 replicon system. Depletion of several RNA-binding or -splicing associated proteins altered viral genomic and subgenomic RNA profiles (Fig. 5a and Extended Data Fig. 4a). Among these candidates, SRSF1 silencing caused one of the strongest reductions in sg.*N/Orf9b* and sg.*Orf7a* RNAs and N protein expression, accompanied by increased g.*Nsp3* RNA and Nsp3 protein levels (Fig. 5a,b and Extended Data Fig. 4a), a pattern resembling that observed with the Rep.KVTA. These results nominated SRSF1 as a candidate host factor that may cooperate with dephosphorylated N to promote subgenomic RNA synthesis.

**Fig. 5.**
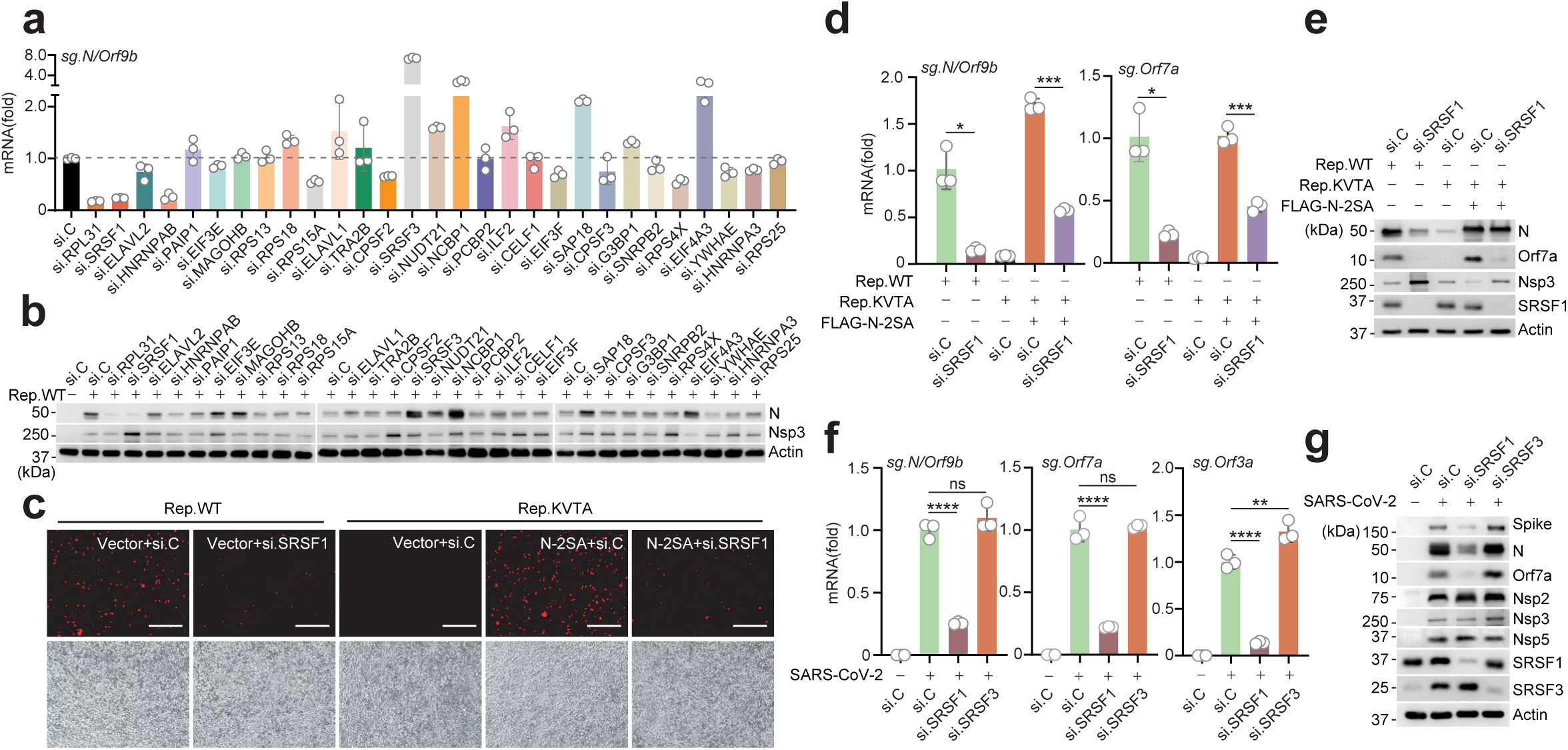
SRSF1 is required for sgRNA synthesis mediated by unphosphorylated N. **a**, RT-qPCR analysis of sg.*N/Orf9b* in HEK293T cells that were transfected for 24 h with the indicated siRNAs and then transfected with Rep.WT for 48 h. Values were normalized to those in si.C-transfected samples, set to 1. **b**, IB analysis of N, Nsp3 and Actin protein expressions from **a**. **c**-**e**, Replication of Rep.WT or Rep.KVTA in HEK293T cells that were transfected for 24 h with either si.C or si.SRSF1 and subsequently transfected for 48 h with Rep.WT or Rep.KVTA together with either empty vector or FLAG-tagged N-2SA, determined by IF microscopy analysis of mScarlet signals (**c**), RT-qPCR analysis of the indicated viral sgRNAs (**d**) and IB analysis of the indicated proteins (**e**). **f**, Viral sgRNA levels in Calu-3 cells that were transfected for 48 h with either si.C, si.SRSF1, or si.SRSF3 and subsequently infected with SARS-CoV-2 (MOI 1) for 12 h, determined by RT-qPCR. **g**, IB analysis of the indicated proteins from the experiment in (**f**). Data are representative of one RNAi screen [conducted in triplicates (**a**)] or at least two independent experiments [mean ± SD of n = 3 biological replicates (**d**,**f**)]. *P < 0.05, **P < 0.01, ***P < 0.001, ****P < 0.0001 (two-tailed Student’s t test with Welch’s correction in **d**, one-way ANOVA with Sidak’s multiple comparison test in **f**). ns, not significant.

Consistent with this proposed concept, SRSF1 depletion markedly reduced the mScarlet signal of Rep.WT (Fig. 5c). Notably, SRSF1 knockdown also impaired the ability of N-2SA to rescue Rep.KVTA (Fig. 5c), as evidenced by reduced N-2SA-mediated restoration of Rep.KVTA-derived sg.*N/ORF9b* and sg.*ORF7a* RNA (Fig. 5d) and ORF7a protein expression (Fig. 5e). By contrast, SRSF1 depletion increased Nsp3 protein expression in both Rep.WT and N-2SA-complemented Rep.KVTA (Fig. 5e). Consistently, in SARS-CoV-2-infected Calu-3 cells, silencing of SRSF1, but not SRSF3, significantly diminished the levels of multiple subgenomic RNAs, including sg.*N/ORF9b*, sg.*ORF7a* and sg.*ORF3a* (Fig. 5f), and decreased expression of the subgenomic RNA-encoded proteins spike, N and ORF7a, whereas the expression of Nsp2, Nsp3 and Nsp5 was minimally affected by SRSF1 depletion (Fig. 5g).

Together, these data identify SRSF1 as an important host factor for SARS-CoV-2 subgenomic RNA synthesis mediated by dephosphorylated N.

### SRSF1 promotes TRS-related RNA binding and higher-order assembly of dephosphorylated N

We next sought to further define the SRSF1-N-interaction. Co-IP analysis confirmed that SRSF1 bound phospho-deficient N-2SA more strongly than to phosphomimetic N-10D (Fig. 6a). Mapping studies using a series of N truncation mutants showed that SRSF1 efficiently associated with full-length N-2SA (N-2SA-FL), whereas deletion of the NTD strongly reduced this interaction. By contrast, the removal of other N domains had either no or minimal effects on SRSF1 binding (Fig. 6a). These data suggest that SRSF1 preferentially binds to the NTD domain of unphosphorylated N.

**Fig. 6.**
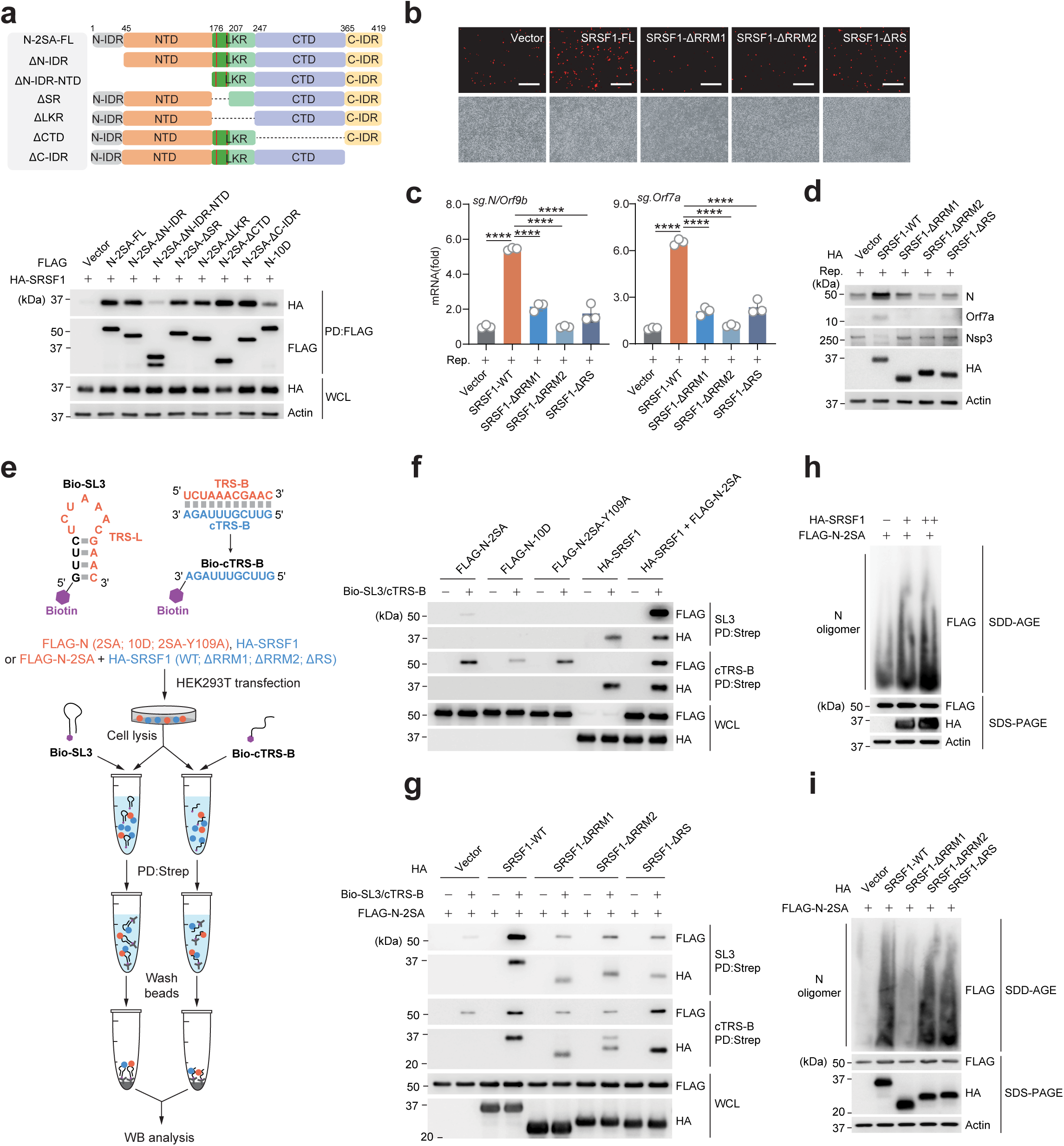
SRSF1 promotes TRS RNA binding and higher-order assembly of dephosphorylated N. **a**, Upper: Schematic representation of SARS-CoV-2 N-2SA full-length (N-2SA FL) or generated truncated mutants. Lower: SRSF1 binding to N in HEK293T cells that were transiently transfected with HA-tagged SRSF1 together with either empty vector or FLAG-tagged N-2SA FL or mutants, determined by PD:FLAG and IB with anti-HA. **b**-**d**, Replication of Rep.WT in HEK293T cells that were co-transfected for 48 h with either empty vector or HA-tagged SRSF1 WT or mutants, determined by IF microscopy analysis of mScarlet signals (**b**), RT-qPCR analysis of the indicated viral sgRNAs (**c**), and IB analysis of the indicated proteins (**d**). **e**, Schematic workflow for the RNA pull-down analysis to assess SRSF1 or N binding to biotinylated SARS-CoV-2 TRS-L (Bio-SL3) or cTRS-B (Bio-cTRS-B). Upper, Generation of Bio-SL3 and Bio-cTRS-B; Lower, HEK293T cells were transfected with either empty vector or the indicated FLAG-tagged N mutants, or HA-tagged SRSF1 WT or mutants, or co-transfected with the indicated combinations. Cell lysates were incubated with the indicated biotinylated RNA probes, followed by streptavidin pull-down (PD:Strep) and IB analysis. **f**, SRSF1 and/or N binding to SARS-CoV-2 Bio-SL3 or Bio-cTRS-B RNA. HEK293T cell lysates expressing the indicated proteins were incubated with Bio-SL3 or Bio-cTRS-B RNA, respectively, followed by PD:Strep and IB analysis with anti-FLAG and anti-HA. **g**, N binding to Bio-SL3 or Bio-cTRS-B RNA under the indicated conditions. HEK293T cell lysates expressing the indicated proteins were incubated with Bio-SL3 or Bio-cTRS-B RNA, respectively, followed by PD:Strep and IB analysis with anti-FLAG and anti-HA. **h, i**, Oligomerization of FLAG-tagged N-2SA in HEK293T cells that were co-transfected for 24 h with HA-tagged SRSF1 WT (**h**) or mutants (**i**), determined by SDD-AGE and IB with anti-FLAG. WCLs were analyzed by SDS-PAGE and probed by IB with anti-FLAG, anti-HA, and anti-Actin (loading control). Data are representative of at least two (**b**-**d** and **f**-**i**) independent experiments. ****P < 0.0001 (one-way ANOVA with Sidak’s multiple comparison test).

We next examined the functional contribution of individual SRSF1 domains to subgenomic RNA synthesis. SRSF1 contains two N-terminal RNA-recognition motifs, RRM1 and RRM2, which mediate RNA binding, followed by a C-terminal arginine/serine-rich RS domain^54,55^. Expression of SRSF1 WT strongly enhanced Rep.WT mScarlet expression, whereas deletion of the RRM1, RRM2, or RS domain significantly compromised this activity (Fig. 6b). Consistently, RT-qPCR and immunoblot analyses showed that while SRSF1 WT effectively increased sg.*N/ORF9b* and sg.*ORF7a* RNA levels as well as ORF7a protein expression, each truncation mutant showed impaired activity (Fig. 6c,d). By contrast, Nsp3 expression was reduced in the presence of SRSF1 WT, but not by any of the truncated mutants (Fig. 6d). Together, this suggests that SRSF1-mediated stimulation of subgenomic RNA synthesis requires its intact domain architecture.

Coronavirus subgenomic RNA synthesis requires discontinuous transcription in which the nascent negative-strand RNA switches template through base-pairing between leader and body transcription-regulatory sequences (TRSs)^31,32^. As N plays a critical role in this process^34–37^, we next asked whether SRSF1 promotes the association of dephosphorylated N with TRS-related RNA elements. To this end, we generated a biotinylated stem-loop containing the leader TRS (Bio-SL3) as well as the complementary body TRS sequence (Bio-cTRS-B) and incubated them with cell lysates containing N and/or SRSF1, followed by streptavidin pull-down (Fig. 6e). Immunoblot analysis showed that N-2SA bound to both SL3 and cTRS-B, whereas the phosphomimetic N-10D showed reduced association with both probes (Fig. 6f), confirming previous findings that N phosphorylation inhibits its RNA-binding ability^13,56^.

Consistent with the role of the NTD in TRS-specific RNA recognition, the RNA-binding-impaired N-2SA Y109A mutant showed markedly reduced binding to Bio-SL3 and more modestly reduced binding to Bio-cTRS-B (Fig. 6f). Notably, SRSF1 associated with both probes and strongly enhanced N-2SA binding to Bio-SL3, with a weaker effect on Bio-cTRS-B. Deletion of RRM1, RRM2 or the RS domain impaired these activities (Fig. 6f,g), indicating that the intact domain architecture of SRSF1 is required for facilitating the binding of dephosphorylated N to TRS-related viral RNA elements.

Given that N higher-order assembly underlies RNA-induced condensate formation and can concentrate viral RNA together with RdRp components to promote coronavirus transcription^16^, we next tested if SRSF1 modulates the assembly state of dephosphorylated N. Semi-denaturing detergent agarose gel electrophoresis (SDD-AGE) analysis showed that WT SRSF1 increased N oligomerization in a dose dependent manner (Fig. 6h). Furthermore, whereas the SRSF1 ΔRRM2 and ΔRS mutants promoted N oligomerization comparably to WT SRSF1, the ΔRRM1 mutant lost this activity (Fig. 6i), indicating that RRM1 is required for SRSF1-driven N higher-order assembly.

Collectively, our data support a model in which SARS-CoV-2 Nsp3, through a conserved PP1-binding motif within the Ubl1 domain, recruits PP1α/γ to dephosphorylate N. Nsp3 promotes the formation of a dephosphorylated pool of N that is competent for RNA binding and functional cooperation with SRSF1. Our mechanistic data revealed that the N-SRSF1 complex facilitates the cooperative engagement of dephosphorylated N with TRS-related viral RNA elements, and also promotes N higher-order assembly, thereby supporting efficient subgenomic RNA synthesis.

## Discussion

The SARS-CoV-2 N is extensively phosphorylated within its serine/arginine-rich linker region, and this modification has emerged as a central mechanism controlling the distinct functions of N during infection. Previous studies have shown that N phosphorylation regulates RNA binding, LLPS, viral replication and genome packaging, supporting the concept that distinct phosphorylation states of N are linked to distinct phases of the coronavirus life cycle^13,14,17,40^. In accord, several host kinases, including SRPK, GSK-3 and CK1 family kinases, have been implicated in the sequential phosphorylation of N, and perturbation of these kinases can impair SARS-CoV-2 replication ^9,11^. However, N phosphorylation inhibits RNA binding and viral RNP assembly, and virion-associated N is thought to be hypophosphorylated or unphosphorylated^10,38,56^. Importantly, how N is dephosphorylated and whether this step determines additional functions of N has remained unclear. Our work identifies Nsp3 as a key orchestrator of N dephosphorylation at the replication organelle. Through its Ubl1 domain, Nsp3 preferentially binds to phosphorylated N and recruits the phosphatases PP1α/γ through a conserved PP1-binding motif, thereby generating a local pool of dephosphorylated N that is needed for RNP assembly. Intriguingly, our data also showed that N dephosphorylation is required for efficient sgRNA synthesis through the recruitment of and functional cooperation with the host splicing factor SRSF1.

Our findings also provide novel insights into Nsp3’s role as a key orchestrator of N function. Nsp3 is a core component of coronavirus replication organelles and, together with Nsp4, forms DMV-associated pore structures proposed to connect the protected replication compartment with the cytosol and permit viral RNA transit^18,57^. Genetic and structural studies have shown that N engages the N-terminal Ubl1-containing region of Nsp3, and that disruption of this interaction impairs coronavirus RNA synthesis, fitness and virulence^4,25,27,58^. The mechanistic role of N’s interaction with Nsp3-Ubl1, however, has been elusive. Our data suggest that Nsp3-Ubl1 is not only a docking site for N but also acts as a scaffold that brings phosphorylated N and host PP1 together at replication organelles. This links N recruitment to the generation of a dephosphorylated N protein pool that binds viral RNA and supports RNP assembly. Consistent with this model, Ubl1 also binds the viral 5′ UTR, and the Ubl1-N complex shows increased affinity for 5′ UTR RNA^29^. Thus, Nsp3 may coordinate viral RNA transit, N dephosphorylation, and N loading onto newly synthesized gRNA at or near the DMV pore. How Nsp3-Ubl1-mediated N dephosphorylation is temporally controlled to balance viral RNA replication at early times of infection vs. genome packaging at late time points warrants further investigation.

Our study also reveals a previously unknown function of N dephosphorylation, specifically its role in supporting efficient viral sgRNA synthesis as indicated by our data using a single-cycle replicon system. Mutation of the conserved PP1-binding motif in Nsp3 selectively abrogated sgRNA production. This places Nsp3-mediated N dephosphorylation and its role in facilitating efficient sgRNA transcription as a critical event that occurs prior to terminal genome packaging. Our findings complement previous work that showed that N supports coronavirus transcription and that phosphorylated N recruits the RNA helicase DDX1 to promote synthesis of gRNA and longer sgRNAs^4,38,46^. Our proteomic analysis further suggests that the phosphorylation state of N partitions it into distinct host protein-binding networks. Phosphorylated N is preferentially associated with ribosome- and translation-related proteins. This may help couple newly synthesized viral RNAs to the translation-associated machinery at replication organelles, consistent with recent evidence that N can organize translation-promoting host factors near DMVs^59,60^. By contrast, dephosphorylated N complexed with several RNA-processing factors, including splicing-related RBPs involved in RNA recognition, RNP assembly and RNA remodeling. Because coronavirus sgRNA synthesis requires long-range template switching during negative-strand synthesis^31,32^, these factors may help shape viral RNA conformations or position dephosphorylated N-TRS for productive leader-body pairing, which warrants future investigation.

SRSF1 is a prototypical SR-family RBP with broad roles in RNA metabolism, including transcriptional control, alternative splicing and mRNA export^61–64^. Interestingly, proteins and certain RNAs encoded by several viruses (i.e., EBV, HPV16 and HIV-1) have been shown to redirect SRSF1/3 activity to regulate the fate of viral RNAs^65–67^. Our data revealed that SRSF1 binds to and cooperates specifically with dephosphorylated N to promote SARS-CoV-2 sgRNA synthesis. Mechanistically, the intact SRSF1 protein domain architecture, including RRM1, RRM2 and RS domains, was required to enhance RNA binding of N and ultimately sgRNA synthesis. By contrast, ability of SRSF1 to promote N oligomerization depended specifically on RRM1. Given that N undergoes RNA-dependent LLPS^16,40,68–70^, this separation of function suggests that SRSF1 may tune this event: the intact RRM1-RRM2-RS architecture of SRSF1 supports productive N loading onto TRS-related RNA elements, whereas RRM1 facilitates N higher-order assembly. Whether SRSF1 directly stabilizes leader-body pairing, remodels local RNA structure, promotes polymerase pausing and/or facilitates annealing during template switching remains to be determined.

Besides SRSF1, our targeted siRNA screen also identified several additional host factors that may contribute to SARS-CoV-2 subgenomic RNA synthesis. In particular, depletion of RPL31 and HNRNPAB also significantly reduced subgenomic RNA levels, suggesting that multiple host RNA-binding or RNA-processing factors may participate in this process. The specific roles of these factors, and whether they function independently of, or cooperatively with, dephosphorylated N, will require further investigation.

Our findings may also have implications for antiviral intervention. N phosphorylation has been proposed as a therapeutic vulnerability, with GSK-3 inhibition impairing N phosphorylation and restricting coronavirus replication^9^. However, GSK-3 has broad cellular functions, and N phosphorylation is controlled by multiple host kinases, including SRPK and CK1 family members^11^. Thus, kinase inhibition may have limited specificity, context-dependent efficacy, and potential toxicity. Our findings extend this concept by revealing the molecular details of the opposing reaction —N dephosphorylation— including the responsible enzymes (PP1α and PP1γ isoforms) and Nsp3’s role in orchestrating this event. Although direct PP1 inhibition would likely be toxic, viral interfaces that position PP1α/γ, including the Nsp3-Ubl1 PP1-docking motif and the Nsp3-N or N-SRSF1 interface, are expected to offer more selective targets and may help develop pan-coronavirus inhibitors.

## Online Methods

### Cells

Human embryonic kidney cells (HEK293T, CRL3216; ATCC) were maintained in Dulbecco’s modified Eagle medium (DMEM, 41965039; Gibco) supplemented with 10% fetal bovine serum (FBS, 10270106; Gibco), 1% penicillin-streptomycin (Pen-Strep, 15140122; Gibco) and 1 mM sodium pyruvate (11360-070; Gibco). Vero E6-TMPRSS2 and A549-ACE2 cells were described previously^43^ and cultured in DMEM supplemented with 10% FBS, 1 mM sodium pyruvate and 1% Pen-Strep. Calu-3 cells (HTB-55; ATCC) were maintained in Eagle’s minimum essential medium (EMEM, 30-2003; ATCC) supplemented with 10% FBS, 1 mM sodium pyruvate and 1% Pen-Strep. All cells were cultured at 37 °C with 5% CO2. All cell lines used in the experiments of our study were regularly tested by PCR assay to validate the absence of mycoplasma contamination.

### Plasmids

Expression plasmids encoding V5-tagged SARS-CoV-2 Nsp3 WT and ΔUbl1 mutant, FLAG-tagged SARS-CoV-2 N WT, N-2SA and N-10D, as well as HA-tagged PP1α, PP1β and PP1γ have been described previously^43,71^. HA-tagged SRSF1 was generated by amplifying human SRSF1 cDNA from a synthetic gBlock gene fragment (purchased from IDT) and subsequent subcloning into the pcDNA3.1(−) vector containing an N-terminal 3×HA tag. The other Nsp3, N, and SRSF1 mutants used in our study were generated by site-directed mutagenesis or overlap-extension PCR. All plasmids were verified by Sanger sequencing (Azenta Life Sciences) or whole plasmid nanopore sequencing (Plasmidsaurus).

### Viruses and replicons

SARS-CoV-2 strain K49 was propagated in Vero E6-TMPRSS2 cells as detailed previously^43^. The SARS-CoV-2 replicon Rep.WT and Rep.KVTA were generated by CPER-based method using a published protocol^43^. Forty-eight hours after transfection, replicon activity was assessed by fluorescence microscopy measuring mScarlet signal, by RT-qPCR analysis of viral genomic and subgenomic RNAs, and/or by immunoblotting of viral proteins. Specifically, mScarlet signals in replicon-transfected HEK293T cells were imaged on an EVOS M5000 Imaging System (Thermo Fisher Scientific) equipped with an RFP (531/593 nm) light cube and a 10× objective. All experiments with infectious SARS-CoV-2, or the replicon, were conducted at the Cleveland Clinic Florida Research and Innovation Center (CC-FRIC) using the appropriate biosafety containments and under protocols approved by the Institutional Biosafety Committee in accordance with NIH guidelines.

### Transfection and siRNA gene knockdown

Transient DNA transfections were performed with linear polyethylenimine (PEI; 1 mg/mL in 10 mM Tris-HCl, pH 6.8), Lipofectamine 2000 (11668027; Thermo Fisher Scientific) or TransIT-X2 Transfection Reagent (MIR 6000; Mirus) according to the manufacturer’s protocols. Unless otherwise stated, HEK293T cells were analyzed at 24 h post plasmid transfection for biochemical assays, and at 48 h after replicon transfection for replication assays. Transient knockdown experiments in A549-hACE2, HEK293T, and Calu-3 cells were performed using siGENOME or ON-TARGETplus SMARTpool small-interfering (si)RNAs (purchased from Horizon Discovery). The specific siRNAs used in this study are: PP1α (M-008927-01), PP1β (M-008685-00), PP1γ (M-006827-00), PP2Aα (M-003598-01), PP2Aβ (M-003599-03), RPL31 (L-013587-00), SRSF1 (L-018672-01), ELAVL2 (L-019801-00), HNRNPAB (L-013483-01), PAIP1 (L-019182-01), EIF3E (L-010518-00), MAGOHB (L-016706-02), RPS13 (L-011150-02), RPS18 (L-011890-01), RPS15A (L-013542-01), TRA2B (L-007278-00), CPSF2 (L-013404-00), SRSF3 (L-030081-00), NUDT21 (L-012335-01), NCBP1 (L-019672-00), PCBP2 (L-012002-00), ILF2 (L-017599-00), CELF1 (L-020166-00), EIF3F (L-019535-00), SAP18 (L-012140-00), CPSF3 (L-006365-01), SNRPB2 (L-016910-01), RPS4X (L-011138-00), EIF4A3 (L-020762-00), YWHAE (L-017302-02), HNRNPA3 (L-019347-00) and RPS25 (L-013629-00).

### RNA isolation and RT-qPCR analysis

Total RNA was isolated from cells using the E.Z.N.A. HP Total RNA Kit according to the manufacturer’s instructions. Equal amounts of RNA were analyzed using the SuperScript III Platinum One-Step RT-qPCR Kit (11732088; Invitrogen) with PrimeTime qPCR Probe Assays (IDT) on a QuantStudio 6 Pro Real-Time PCR System (Applied Biosystems). Viral genomic RNA was measured using probes targeting g.*Nsp3*, and viral subgenomic RNAs were measured using probes targeting sg.*N/Orf9b*, sg.*Orf7a* or sg.*Orf3a*, using methods previously described^72^. Relative mRNA expression was normalized to the levels of *HPRT1* (cellular housekeeping gene) and expressed relative to the values for control cells using the ΔΔ*C*_t_ method.

### Cell lysis and immunoprecipitation

Cells that were transfected as indicated (see details in figure legends) were washed with PBS and lysed in NP-40 buffer (50 mM HEPES, pH 7.4, 150 mM NaCl, 1% NP-40, 1 mM EDTA and 1x protease inhibitor cocktail). Lysates were cleared by centrifugation at 21,000 × *g* for 15 min at 4 °C. For FLAG- or HA-tagged-protein pulldown (PD) assays, cleared lysates were incubated with anti-FLAG M2 (M8823; Sigma-Aldrich) or anti-HA (88837; Thermo Fisher Scientific) magnetic beads for 4–16 h at 4 °C. For immunoprecipitation (IP) with anti-V5, cell lysates were incubated with Protein G Dynabeads (10004D; Thermo Fisher Scientific) conjugated with the anti-V5 antibody (R960-25; Invitrogen) at 4°C for 4–16 h. For endogenous Nsp3 IP, lysates were incubated with Protein G Dynabeads conjugated with the anti-Nsp3 antibody (GTX135589; GeneTex) at 4°C for 4–16 h. Normal Rabbit IgG (2729S; CST)- conjugated beads served as the specificity control where indicated. Beads were washed three to five times with the corresponding lysis buffer and bound proteins were released by heating in 1x Laemmli SDS sample buffer (S3401; MilliporeSigma) at 95 °C for 5 min.

### Immunoblotting and antibodies

Immunoprecipitated protein samples or WCLs were resolved by 8–12% Bis-Tris SDS-PAGE and transferred onto PVDF membranes (Cat# 1620177; Bio-Rad), followed by visualization of the blot using the SuperSignal West Pico PLUS or Femto chemiluminescence reagents (Thermo Fisher Scientific) on an ImageQuant LAS 4000 Chemiluminescent Image Analyzer (General Electric) as previously described^43^. The following primary antibodies were used in this study: anti-V5 (1:1,000, R960-25; Invitrogen; used for IP), anti-V5 (1:1,000, 13202; CST; used for immunoblotting of IP and WCLs), anti-Actin (1:2,000, GTX629630; GeneTex), anti-GAPDH (1:1,000, 97166; CST), anti-FLAG Rabbit mAb (1:1,000, 14793; CST), anti-HA (1:1,000, 3724; CST), anti-FLAG M2 Mouse mAb (1:2,000, F1804; MilliporeSigma), anti-PP1α (1:2,000, A300-904A; Thermo Fisher Scientific), anti-PP1β (1:2,000, A300-905A; Thermo Fisher Scientific), anti-PP1γ (1:2,000, A300-906A; Thermo Fisher Scientific), anti-PP2Aα (1:1,000, 13482-1-AP; Thermo Fisher Scientific), anti-PP2Aβ (1:1,000, 12554-2-AP; Thermo Fisher Scientific), anti-NSP3 (1:1,000, GTX135589; GeneTex), anti-Nucleocapsid (N) (1:1,000, 33717; CST), anti-Nsp2 (1:1,000, 40485S; CST), anti-Nsp5 (1:1,000, 51661S; CST), anti-Spike (1:1,000, 56996S; CST), anti-Orf7a (1:1,000, 67750S; CST), anti-SRSF1 (1:1,000, sc-33652; Santa Cruz) and anti-SRSF3 (1: 1,000, sc-13510; Santa Cruz). Secondary (HRP-linked) antibodies that were used for IB analyses are: anti-mouse IgG (1:5000, 7076; CST) and anti-rabbit IgG (1:5000, 7074; CST).

### RNA pulldown assays

Biotinylated SARS-CoV-2 RNA probes corresponding to the leader TRS-containing SL3 element (Bio-SL3; 5′-biotin-GUUCUCUAAACGAAC-3′) and complementary body TRS sequence (Bio-cTRS-B; 5′-GUUCGUUUAGA-3′-biotin) were custom-synthesized by IDT. HEK293T cells were transfected with empty vector, FLAG-tagged N constructs (i.e., WT and mutants) and/or HA-tagged SRSF1 constructs (i.e., WT and mutants) as indicated. At 24 h after transfection, cells were lysed in RNA pulldown buffer (50 mM HEPES, pH 7.4, 150 mM NaCl, 1% NP-40, 1 mM EDTA, 1 U/µL RNase inhibitor and 1 x protease inhibitor cocktail (P2714; MilliporeSigma)), and lysates were cleared by centrifugation at 21,000 × *g* for 15 min at 4 °C. Equal cell lysates aliquots were incubated with 0.1 µM Bio-SL3 or Bio-cTRS-B RNA probe for 1 h at 4 °C. RNA-protein complexes were then captured using streptavidin-conjugated magnetic beads (11205D; Thermo Fisher Scientific) for an additional 1 h at 4 °C. Afterwards, beads were washed four times with RNA pulldown buffer with vortexing briefly in-between washes. Bound proteins were eluted in 1 x Laemmli SDS sample buffer (S3401; MilliporeSigma). Samples were analyzed by immunoblotting with anti-FLAG and anti-HA antibodies.

### Purification of N-2SA and N-10D-interacting proteins and comparative interactome proteomics analysis

To compare the protein interactomes associated with phospho-deficient vs. phosphomimetic N protein, HEK293T cells (∼4 × 10⁷ cells per group) were transfected for 48 h with 40 µg of FLAG empty vector or FLAG-tagged N-2SA or N-10D. Cells were washed with PBS once and lysed in 1% NP-40 lysis buffer. Lysates were then clarified by centrifugation at 21,000 × *g* for 15 min at 4 °C and incubated with anti-FLAG M2 magnetic beads for 16 h at 4 °C. Beads were washed stringently with NP-40 lysis buffer, and bound proteins were eluted in 1× Laemmli SDS sample buffer. Samples were separated on 4–12% Bis-Tris gels (NP0335BOX; Thermo Fisher Scientific) and proteins visualized by silver staining. Gel lanes were excised, subjected to in-gel trypsin digestion and analyzed by LC-MS/MS at the Cleveland Clinic Research Proteomics Core (Ohio). The LFQ intensities, normalized to the total amount of peptides in each sample (MaxLFQ), were used to compare relative protein abundances across samples. After excluding proteins that were also present in the vector control, proteins detected in the N-2SA or N-10D interactomes were subjected to further analysis. Among these, proteins with MaxLFQ_N-10D/MaxLFQ_N-2SA > 1 were classified as preferentially associated with N-10D (total of 176 proteins), whereas proteins with MaxLFQ_N-10D/MaxLFQ_N-2SA < 1 were classified as preferentially associated with N-2SA (total of 146 proteins). These two protein sets were used for Gene Ontology-Biological Process (GO-BP) and KEGG pathway enrichment analyses. For PPI network visualization, proteins with |log₂(MaxLFQ_N-10D/MaxLFQ_N-2SA)| > 1 were highlighted, and the relative protein abundance within each group was represented by node size based on spectral abundance factor (SAF) values.

### Gene Ontology enrichment, KEGG, and PPI network analyses

GO-BP and KEGG pathway enrichment analyses were performed using DAVID Bioinformatics Resources v6.8. Specifically, two protein sets were analyzed: proteins preferentially associated with N-2SA (146 proteins) and proteins preferentially associated with N-10D (176 proteins). Protein identifiers were converted to official human gene symbols and analyzed using *Homo sapiens* as the reference background. Enrichment significance was determined using DAVID’s modified Fisher’s exact test (EASE score), and terms were ranked by false discovery rate (FDR)-adjusted P values. Redundant or closely related GO terms were consolidated for visualization, and representative terms were selected based on statistical significance and biological relevance. In dot plots, color indicates the FDR-adjusted P value, and dot size indicates fold enrichment relative to the whole-genome background. PPI networks for the N-2SA and N-10D interactomes were retrieved from the STRING database and visualized using Cytoscape (version v3.10.4). Edges represent known or predicted protein-protein associations from STRING, and node color gradient reflects log₂(MaxLFQ_N-10D/MaxLFQ_N-2SA), and node size indicates relative protein abundance based on SAF values.

### N oligomerization assay using SDD-AGE

N oligomerization was examined by SDD-AGE following specific procedures previously described^43^. Briefly, FLAG-tagged N-2SA was transfected into HEK293T cells together with either empty vector or HA-tagged SRSF1 WT or mutants. Twenty-four hours later, cells were lysed in a buffer containing 50 mM HEPES (pH 7.4), 150 mM NaCl, 0.5% (v/v) NP-40, 10% (v/v) glycerol and 1× protease inhibitor cocktail (MilliporeSigma) at 4°C for 20 min. Cell lysates were clarified by centrifugation at 16,000 × *g* for 10 min at 4°C and then incubated on ice for 1 h. Cell lysates were then mixed with 4× SDD-AGE sample buffer (2× Tris/borate/EDTA (TBE), 40% (v/v) glycerol, and 8% (w/v) SDS) to a final concentration of 1× (0.5× TBE, 10% (v/v) glycerol, and 2% (w/v) SDS), and then incubated for 5 min at room temperature. Samples were then resolved on a vertical 1.5% agarose gel containing 1× TBE and 0.1% (w/v) SDS at 100V for 40 min at 4°C. Proteins were transferred onto a PVDF membrane and analyzed by IB with anti-FLAG.

### Sequence alignment and structural modelling

Nsp3-Ubl1 sequences from representative alpha-, beta-, and gamma-coronaviruses were aligned with MUSCLE^73^, and the conserved PP1-binding motif K/R(X)VXF was highlighted. The solved structure of Nsp3-Ubl1 (7KAG) was accessed from the RCSB Protein Data Bank^74^. Predicted complexes containing SARS-CoV-2 Nsp3-Ubl1 (residues 1–111), human PP1γ (residues 1–300) and the SARS-CoV-2 N linker region (N-LKR; residues 180–256) were modeled using AlphaFold3^75^. For phosphorylated N-LKR modelling, selected serine and threonine residues within the SR region were specified as phosphoserine and phosphothreonine, respectively, using the AlphaFold3 post-translational modifications (PTMs) module. Models with the highest pTM and ipTM scores, with both scores >0.6, were selected for visualization. Structural models were visualized and rendered using PyMOL v.2.5.5.

### Statistical analysis

Data were analyzed using the GraphPad Prism v.11 software. Unless otherwise stated, comparisons between two groups were performed using two-tailed unpaired Student’s t-tests, and comparisons among multiple groups were performed using one-way ANOVA with Sidak’s multiple-comparisons test. P values of < 0.05 were considered statistically significant. Significance is indicated as *P < 0.05, **P < 0.01, ***P < 0.001 and ****P < 0.0001. No pre-specified effect sizes were assumed, and the number of replicates is specified in the corresponding figure legends.

## Data availability

The data that support the findings of this study are available from the corresponding author upon request. The raw mass spectrometry data were submitted to PRIDE Archive (Project accession: PXD080804).

## Acknowledgements

We are grateful to Belinda Willard (Proteomics Core, Cleveland Clinic Research, Ohio) for support with the mass spectrometry analysis. Furthermore, we thank previous Gack lab members, specifically GuanQun Liu (now McGill University), for his technical support with constructing the replicon. This study was supported by grants from the US National Institutes of Health (AI087846 and AI169444 to M.U.G.).

## Author contributions

J.Z. and M.U.G. conceptualized the study. J.Z. designed, performed and analyzed all experiments of this study. J.Z., C.C., and M.U.G. wrote and edited the manuscript. M.U.G. supervised the study and acquired funding.

## Competing interests

The authors declare no competing interests.

**Supplementary Figure 1. Analysis of Nsp3, N and PP1 complex formation.** (a) Modeling of the Nsp3 Ubl1, unphosphorylated N-LKR, and PP1γ complex formation using AlphaFold3 (pTM: 0.77; ipTM: 0.74). The PP1-binding pocket, catalytic site, and Ubl1&LKR binding site are highlighted. (b) Superimposition of unphosphorylated N-LKR (pink) and phospho-simulated N-LKR (purple) in complex with Nsp3 Ubl1 and PP1γ, corresponding to (a) and Fig. 2h. The PP1-binding pocket, catalytic site, and Ubl1&LKR binding site are highlighted. (c) Nsp3-N binding in HEK293T cells that were transfected for 24 h with either empty vector (−) or FLAG-tagged N together with V5-tagged Nsp3 WT, AVAA or ΔUbl1, determined by PD:FLAG and IB with anti-V5. Data are representative of at least two (c) independent experiments.

**Supplementary Figure 2. Functional characterization of the SARS-CoV-2 replicon encoding Nsp3-KVTA.** (a) PP1γ-Nsp3 binding in HEK293T cells that were transfected for 24 h with either empty vector (−) or HA-tagged PP1γ together with V5-tagged Nsp3 WT, AVAA or KVTA, determined by IP:HA and IB with anti-V5. (b) IB analysis of FLAG-tagged N WT or S2A in HEK293T cells that were co-transfected for 24 h with V5-tagged Nsp3 WT or the mutants AVAA or KVTA. (c) RT-qPCR analysis of the indicated viral RNAs in HEK293T cells that were transfected for 48 h with either Rep.WT or Rep.KVTA. (d) IF microscopy analysis of mScarlet signals in HEK293T cells that were either mock-treated or transfected for 48 h with either Rep.WT and empty vector, or with Rep.KVTA together with empty vector or the indicated plasmids. (e) IB analysis of the indicated protein expression from (d). (f) IF microscopy analysis of mScarlet signals in HEK293T cells that were transfected for 48 h with either Rep.WT and empty vector, or with Rep.KVTA together with either empty vector or FLAG-tagged N-10D or N-2SA. (g) IB analysis of the indicated viral proteins from (f). (h) IB analysis of the indicated viral proteins in HEK293T cells that were transfected for 48 h with either Rep.WT and empty vector, or with Rep.KVTA and empty vector or FLAG-tagged N-S2A or N-2SA-Y109A. Data are representative of at least two independent experiments. **P < 0.01, ***P < 0.001 (two-tailed Student’s t test with Welch’s correction).

**Supplementary Figure 3. Protein-protein interaction (PPI) networks of unphosphorylated and phosphorylated N protein.** (a, b) PPI network visualizations of the N-2SA and N-10D interactomes generated using Cytoscape. The color gradient indicates log_2_ fold change in MaxLFQ intensity, calculated as Log₂[MaxLFQ_N-10D/MaxLFQ_N-2SA]. Negative values are shown in blue and denote proteins preferentially associated with N-2SA, whereas positive values are shown in red and denote proteins preferentially associated with N-10D Proteins with |log₂FC| > 1 are highlighted. Dot size represents relative abundance of the protein based on SAF values. Data shown are from one unbiased AP-MS screen.

**Supplementary Figure 4. A targeted siRNA screen of host factors required for viral subgenomic RNA synthesis.** RT-qPCR analysis of sg.*Orf7a* (upper) and g*.Nsp3* (lower) in HEK293T cells that were transfected for 24 h with the indicated siRNAs and then transfected with SARS-CoV-2 replicon (WT) for 48 h. si.C, control nontargeting siRNA. Data are representative of one RNAi screen (conducted in triplicates).

## Notes

### Competing Interest Statement

The authors have declared no competing interest.

