## Supplementary figures and images for "Nsp3 Ubl1-orchestrated dephosphorylation of N protein promotes coronaviral subgenomic RNA synthesis"

### Supplementary Figures 1-4

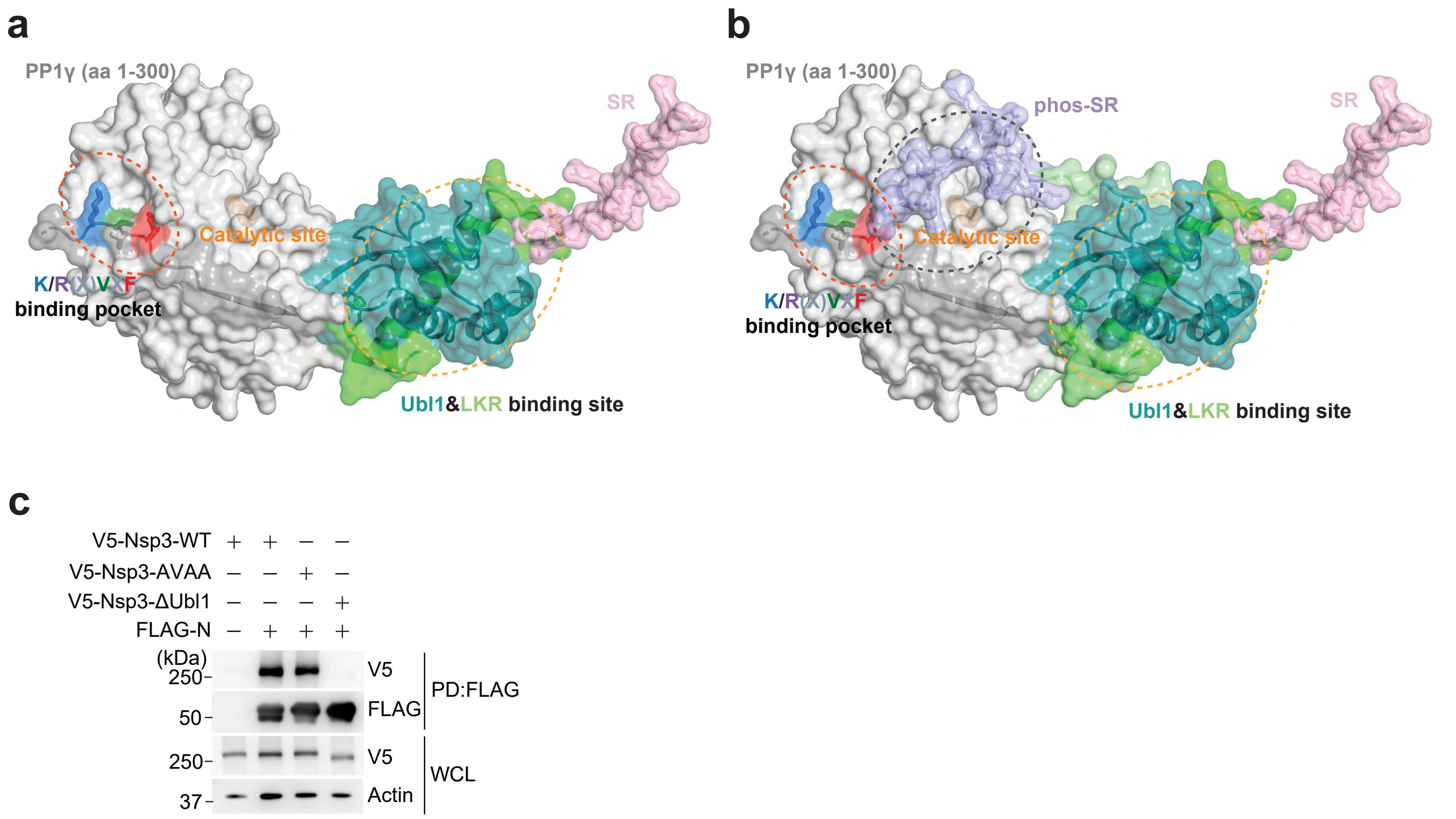

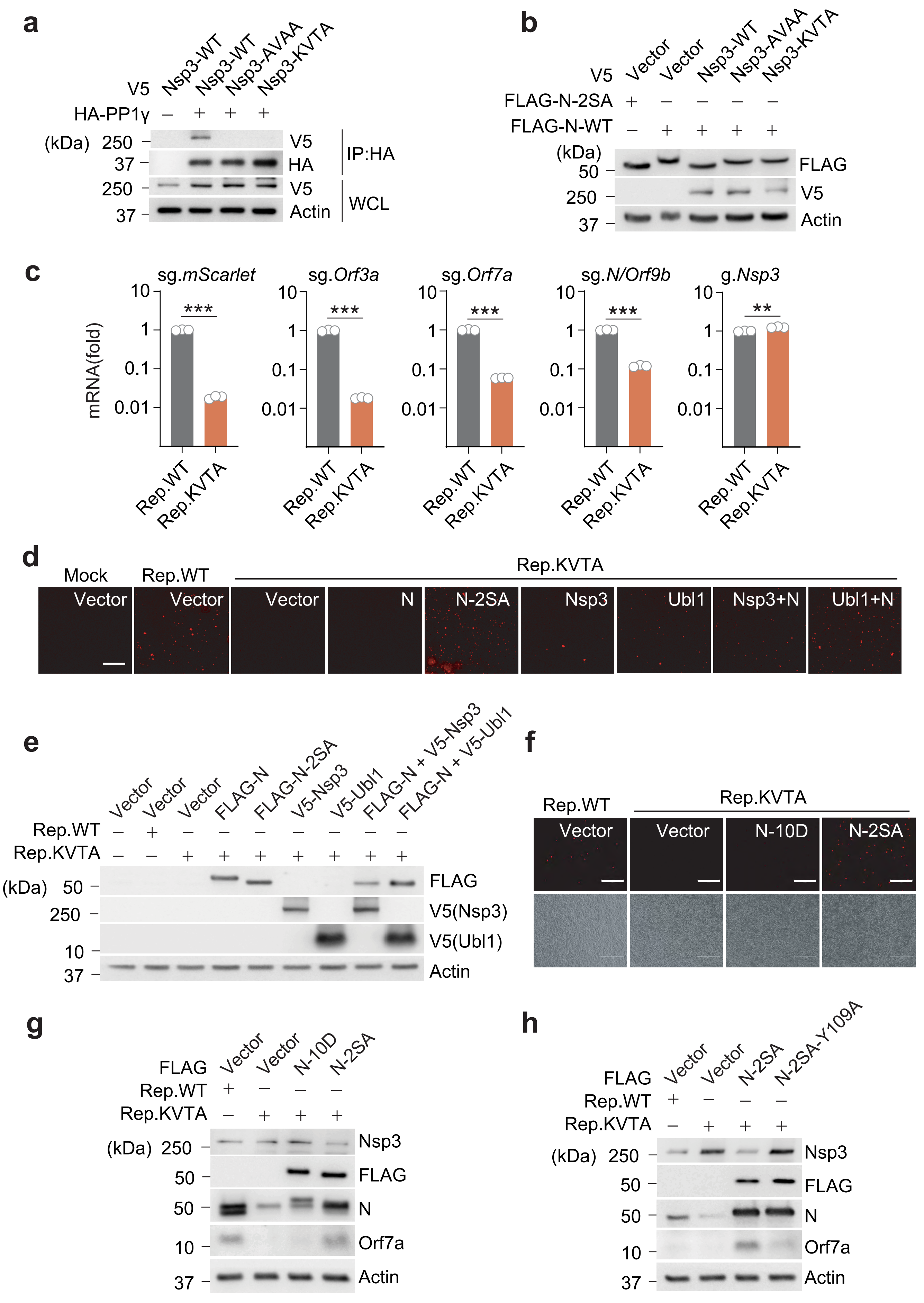

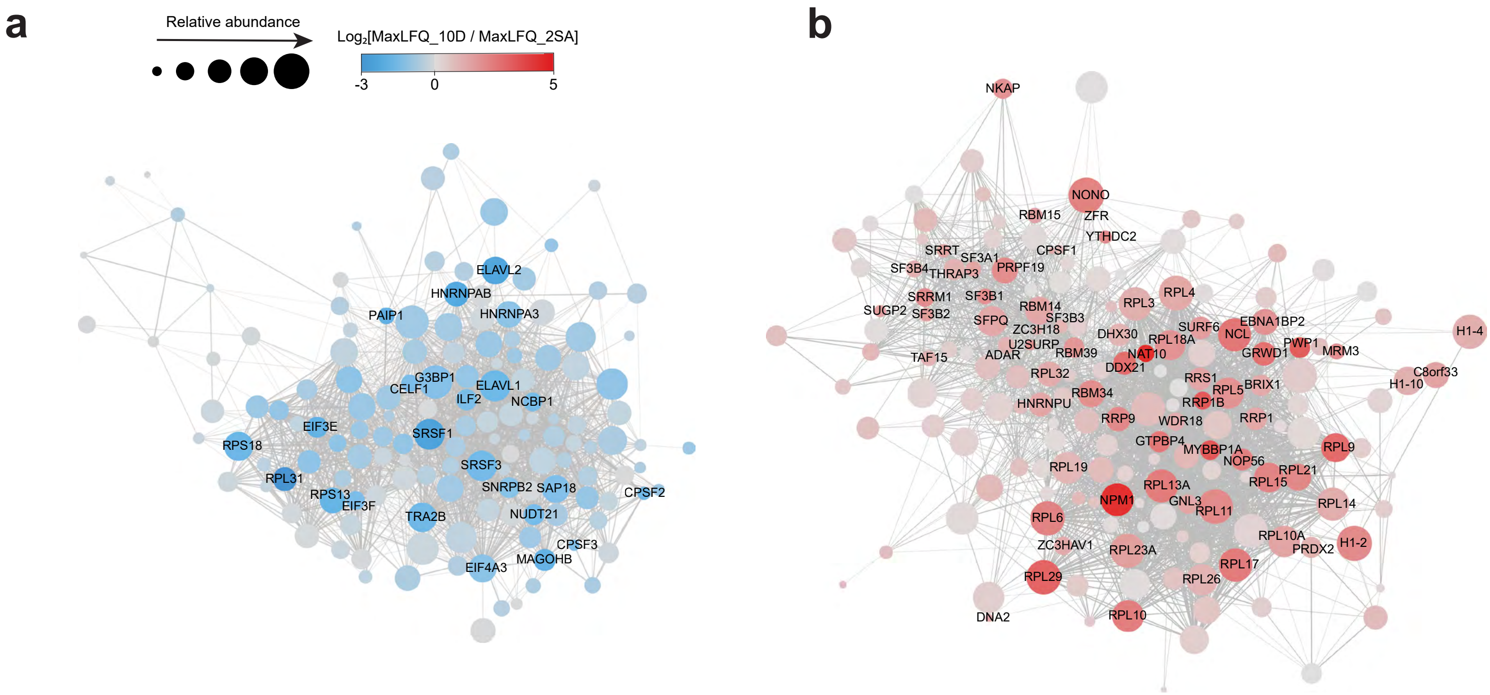

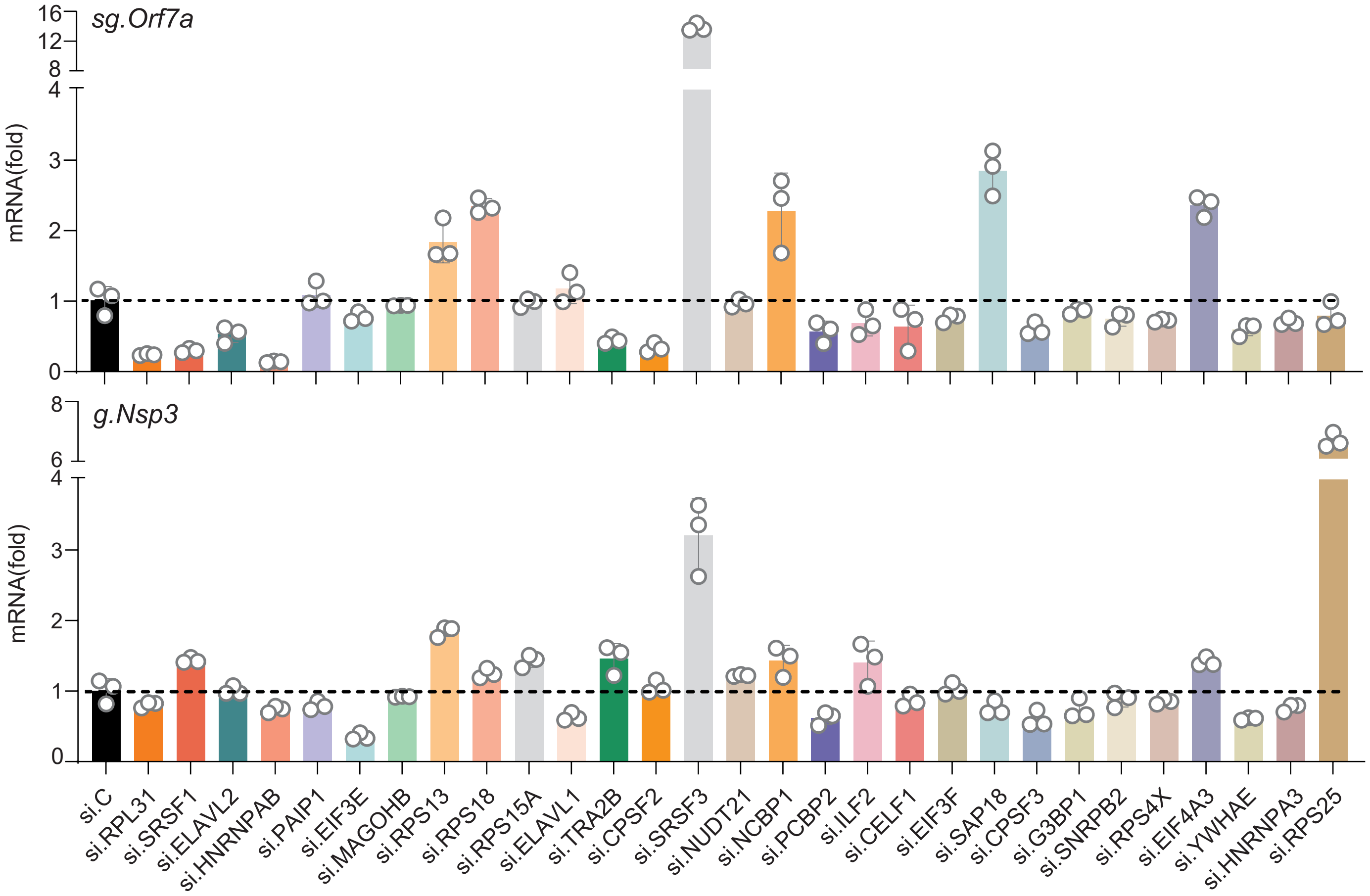
